# Yeast-to-hypha transition of *Schizosaccharomyces japonicus* in response to natural stimuli

**DOI:** 10.1101/481853

**Authors:** Cassandre Kinnaer, Omaya Dudin, Sophie G Martin

## Abstract

Many fungal species are dimorphic, exhibiting both unicellular yeast-like and filamentous forms. *Schizosaccharomyces japonicus*, a member of the fission yeast clade, is one such dimorphic fungus. Here, we first identify fruit extracts as natural, stress-free, starvation-independent inducers of filamentation, which we use to describe the properties of the dimorphic switch. During the yeast-to-hypha transition, the cell evolves from a bipolar to a unipolar system with 10-fold accelerated polarized growth but constant width, vacuoles segregated to the non-growing half of the cell, and hyper-lengthening of the cell. We demonstrate unusual features of *S. japonicus* hyphae: these cells lack a Spitzenkörper, a vesicle distribution center at the hyphal tip, but display more rapid cytoskeleton-based transport than the yeast form, with actin cables being essential for the transition. *S. japonicus* hyphae also remain mononuclear and undergo complete cell divisions, which are highly asymmetric: one daughter cell inherits the vacuole, the other the growing tip. We show these elongated cells scale their nuclear size, spindle length and elongation rates but display altered division size controls. This establishes *S. japonicus* as a unique system that switches between symmetric and asymmetric modes of growth and division.

## Introduction

Cellular morphologies are extremely varied. However, the overall mechanisms generating polarity are thought to be conserved across the species (Nelson, 2003). In fungi, whose shapes are defined by an external rigid cell wall, the location of polarity factors on specific cortical regions locally drives cell growth through cell wall expansion and remodeling, to generate specific cell morphologies. Many fungal species are dimorphic, exhibiting distinct morphologies depending on growth conditions. In this study, we used the fission yeast *Schizosaccharomyces japonicus* (*S. japonicus*), a dimorphic species from the early diverging ascomycete fission yeast clade, to describe the changes occurring during the dimorphic switch.

*S. japonicus* is estimated to have diverged 220Mya from its well-studied cousin *Schizosaccharomyces pombe* (*S. pombe*), with which it displays at least 85% orthologous genes (Rhind et al., 2011). It can grow either in the yeast form, of dimensions slightly larger than *S. pombe*, or in a filamentous form (Niki, 2014). While fungal dimorphism is usually associated with pathogenicity (Nemecek et al., 2006), *S. japonicus* is non-pathogenic to humans making it a convenient model to study the transition in growth mode. It was initially isolated on strawberries from a field in Japan in 1928 (Yukawa and Maki, 1931) and a variant was discovered over a decade later in grape extracts by an American team (Wickerham and Duprat, 1945). The *S. japonicus* yeast form resembles *S. pombe*: cells are rod-shaped, divide medially, grow in a bipolar manner (Sipiczki et al., 1998a) and use the small GTPase Cdc42 for cell morphology (Nozaki et al., 2018). In *S. pombe*, Cdc42 controls cell shape by activating the formin For3 and the exocyst complex for polarized exocytosis of secretory vesicles (Martin and Arkowitz, 2014). However, it also displays important differences, notably in having a semi-open mitosis (Yam et al., 2011) and in division site positioning. In *S. pombe*, septum positioning relies on positive signals from the nucleus, itself placed medially by associated microtubules pushing against both cell poles, and on negative signals preventing septum assembly at cell poles. The anillin-related protein Mid1 conveys the positive signal, whereas the DYRK-family kinase Pom1 serves to inhibit septation at cell poles (Celton-Morizur et al., 2006; Chang et al., 1997; Huang et al., 2007; Padte et al., 2006; Sohrmann et al., 1996). In *S. japonicus*, Pom1 kinase similarly controls medial division, but Mid1 is not required for division site placement (Gu et al., 2015).

*S. japonicus* filamentous form is triggered in response to environmental stresses (Sipiczki et al., 1998b), such as nutritional or nitrogen starvation, and DNA damage stresses (Furuya and Niki, 2010), suggesting that the switch from a small cell to a fast growing hypha serves as an escape mechanism from harsh environmental conditions. Filamentous growth is also light-repressed, as blue light perception by two *white-collar* light receptors present in *S. japonicus* and not in *S. pombe* induces hyphal cell division (Okamoto et al., 2013). Filamentous growth in *S. japonicus* is poorly characterized, though it is thought to share some traits common to other filamentous fungi, such as the presence of a large vacuole at the back of the cell (Sipiczki et al., 1998a).

Filamentous fungi, whether dimorphic (such as *Candida albicans* or *Ustilago maydis*) or not (like *Neurospora crassa* or *Aspergillus nidulans*) grow through rapid apical extension mediated by a vesicle flux towards the growing tip (Riquelme, 2013). Polarized trafficking of vesicles provides the necessary membrane and wall-remodeling material to accommodate the rapid growth of the filamentous form. Vesicles targeted for tip fusion typically accumulate in a spherical organelle, called the Spitzenkörper, which is located close to the growing tip and controls hyphal growth rate and orientation (Riquelme and Sanchez-Leon, 2014). Lower fungi, such as Zygomycetes, and non-fungal Oomycetes do not require a Spitzenkörper to grow but most other filamentous fungi assemble one and it is generally described as a landmark of true filamentous growth (Grove and Bracker, 1970; Read et al., 2010). The hyphal form of filamentous fungi and dimorphic yeasts is multinuclear and its cytoplasm can be compartmentalized by septa that may be incomplete, maintaining cytosolic connection (reviewed in (Steinberg et al., 2017)). This contrasts with the yeast form, which is generally mononuclear and undergoes complete septal division, underlying a need for proper spatial coordination between mitosis and cytokinesis.

In this work, we describe the *S. japonicus* switch from yeast to hypha. We first identify fruit extracts as new inducers of hyphal formation that are independent of nutrient starvation. The *S. japonicus* hyphal form grows much faster and longer than the yeast form, but displays unique features amongst filamentous fungi. Indeed, it lacks a Spitzenkörper, undergoes complete cell divisions and remains mononuclear. We find that cytoskeleton-based transport is more rapid in the hyphal than yeast form, with actin cables necessary for polarized growth, while microtubules contribute to nuclear positioning. *S. japonicus* hyphae divide asymmetrically: the front cell inherits a larger portion of the cytoplasm and no vacuole, and exhibit altered size, growth and division controls. Thus, the *S. japonicus* yeast-to-hypha transition involves the conversion of a symmetric to an asymmetric cell.

## Results

### Fruit extracts induce filamentation in *Schizosaccharomyces japonicus*

Since *S. japonicus* was originally isolated from strawberries and grapes (Wickerham and Duprat, 1945; Yukawa and Maki, 1931), which may represent a natural habitat, we investigated whether these fruits alter the fungus growth behavior. Previous work established that induction of *S. japonicus* filamentation occurs upon stress by nutrient depletion and/or DNA damage (Aoki et al., 2017). On solid rich media in absence of stress, *S. japonicus* primarily grows in the yeast form (Fig. 1A). By contrast, within 3 days of growth on rich media plates supplemented with fruit extracts, *S. japonicus* colonies extended filaments at their periphery, appearing as a white halo around the yeast colony. The filamentation observed at colony edges was invasive as it persisted after plate washing, indicating that the elongated cells have penetrated the solid media (Fig. 1A). Invasive growth was observed with grape (red or white) and strawberry extracts, but also with other berry extracts. Filamentation was increased in presence of higher concentration of red grape extract and decreased with lower concentrations (Fig. 1B). In this work we used 10% red grape extract (RGE) to induce filamentation. RGE did not induce filamentation in other fission yeast species, nor in *Saccharomyces cerevisiae*, which can form pseudohyphae in certain conditions (Gimeno et al., 1992) (Fig. 1C). We note that the ability of RGE to induce filamentation on rich media suggests this is independent of nutrient stress, contrasting with previous reports associating filamentation with escape from stress (Furuya and Niki, 2010; Sipiczki et al., 1998b). This underlies the existence of different triggers and/or mechanisms by which *S. japonicus* transitions in growth forms. A tropism assay showed that *S. japonicus* filaments formed at least as much towards the red grape extract as away from it (Fig. 1D-E). Thus, although we cannot fully exclude oxidative stress as the trigger for the fruit extract-induced switch, this indicates it is not a repellent. Initial characterization of the molecular properties of the RGE inducer showed that it is unlikely to be a nucleic acid, a protein or a lipid and that it is heat-resistant. Phase separation with chloroform/methanol further defined that the inducer is water-soluble. However, the molecular identity of the inducer remains to be identified, as limited screening through candidate molecules present in fruit extracts was so far unsuccessful (Table 1). In summary, fruit extracts represent new, likely stress-free, inducers for the switch to hyphal growth in *S. japonicus*.

Table 1: Hyphal inducing properties of fruit extracts and candidate molecules

**Figure 1.**
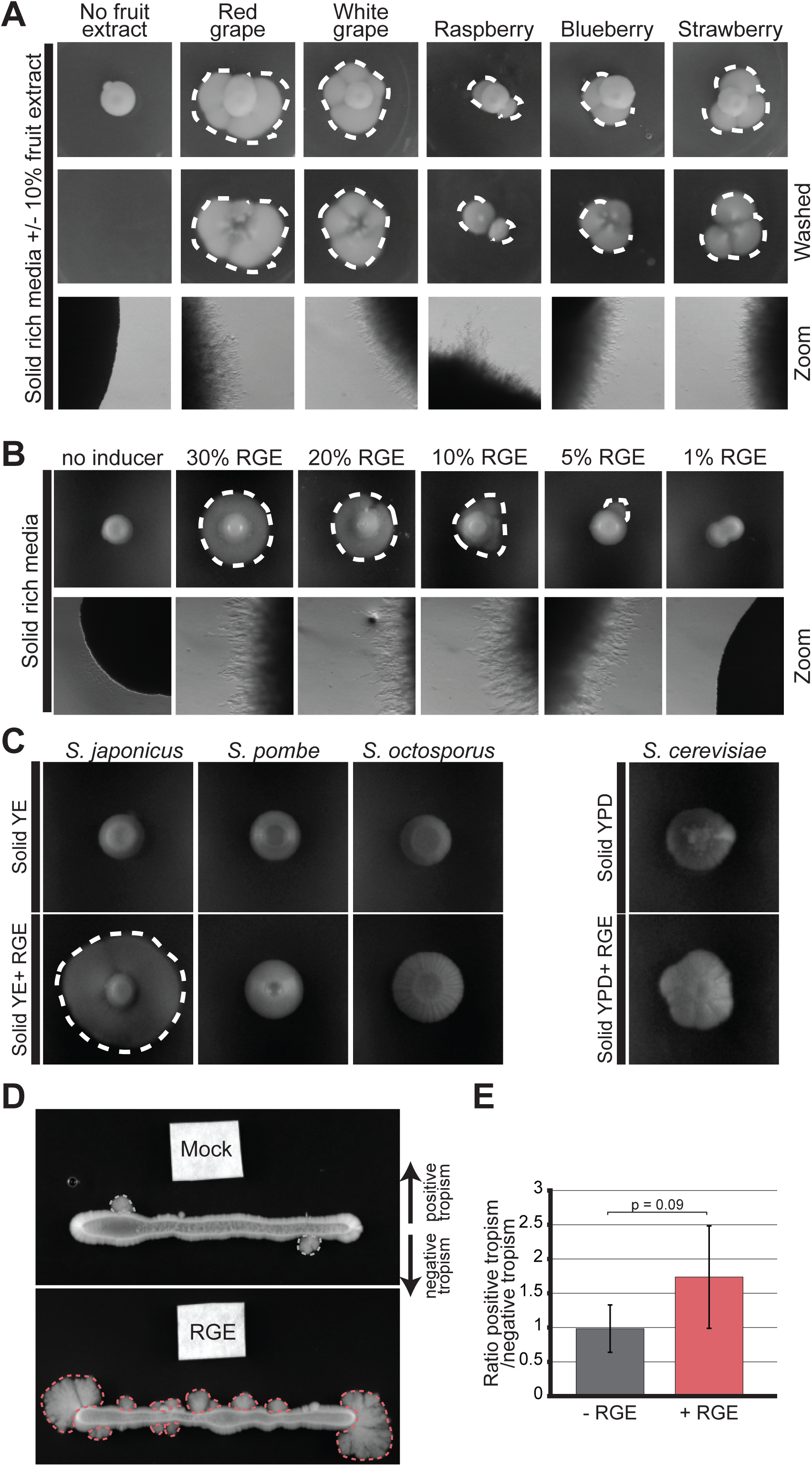
Fruit extracts induce invasive filamentation in *S*. ***japonicus***. **A.** *S. japonicus* growing in agar plates on solid rich media (YE), supplemented or not with 10% fruit extracts before (up) and after (middle) washing of the plate, as well as the same plates imaged under a stereomicroscope (down). **B.** *S. japonicus* growing on solid rich media supplemented or not with a range of concentration of red grape extract (RGE) and the same plate under the stereomicroscope. **C.** *S. japonicus*, *S. pombe*, *S. octosporus* and *S. cerevisiae* growing on solid rich media (YE or YPD) and supplemented or not with RGE. **D.** Tropism experiment to assess the directionality of *S. japonicus* hyphal growth. White filter squares were soaked with YE (up) or RGE (down). **E.** Ratio of positive vs. negative growth tropism in experiments such as in (D). Positive tropism denotes growth towards and negative tropism away from the filter; P = 0.09, t. test. Error bars show standard deviations. Dotted lines highlight penetrative filamentous growth.

### The yeast-to-hypha transition involves extreme vacuolization and dramatic increase in cell size

Microscopy of hyphal cells growing on solid media proved to be challenging due to the invasiveness of hyphae. Therefore, we performed imaging experiments in microfluidic chambers. In this set up, the cells are trapped between a flexible top layer made out of polydimethylsiloxane and a bottom glass layer, neither of which they can penetrate. Because blue light is inhibitory to filamentation (Okamoto et al., 2013), long-term microscopy was performed either with cells carrying a deletion of the white-collar light receptors Wsc1 and Wsc2 or in the presence of a blue-light filter. In these growth conditions, we observed a progressive transition over 24h to the filamentous form at the edges of micro-colonies (Fig. 2A) (Movie S1). Three successive stages in the transition from yeast to hypha can be described. The first landmark of filamentation is the apparition of multiple vacuoles all over the cytoplasm (Sipiczki et al., 1998a), forming a vacuolated yeast form. The vacuoles then polarized to one cell end in what we will refer as the transition form. Finally, once the vacuoles fused together into one large vacuole we refer to them as the hyphal form (Fig. 2B). While the yeast and vacuolated yeast forms mainly grow in a bipolar manner, the transition and hyphal forms are always monopolar underlying a change in mode of growth (Fig. 2C). In time course experiments, the earliest sign of vacuolization was observed 12h after RGE induction, vacuole polarization at one end of the cell occurred after 18h and hyphae were observed 24 hours after induction (Fig. 2D). In the hyphal form, recognizable by its partitioning of the cytosol to the growing end of the cell and the vacuole to the back end, cell extension was strongly correlated with vacuole growth, suggesting that turgor pressure is an important driving force for growth (Fig. 2E). Hyphae also grow very fast, about ten times faster than the yeast form (Fig. 2F). Indeed, hyphae extend at an average rate of 0.58 µm/min, whether RGE is added to rich or minimum medium, a rate comparable to that observed in other filamentous fungi, such as *Candida albicans* and *Aspergillus nidulans* which respectively grow at an average rate of 0.76 µm/min and 0.5 µm/min (Gow and Gooday, 1982; Horio and Oakley, 2005). The monopolar transition form also displayed rapid growth rates, though slightly slower than hyphae, suggesting that transition to monopolar growth and growth rate increase are simultaneous events (Fig. 2F).

**Figure 2.**
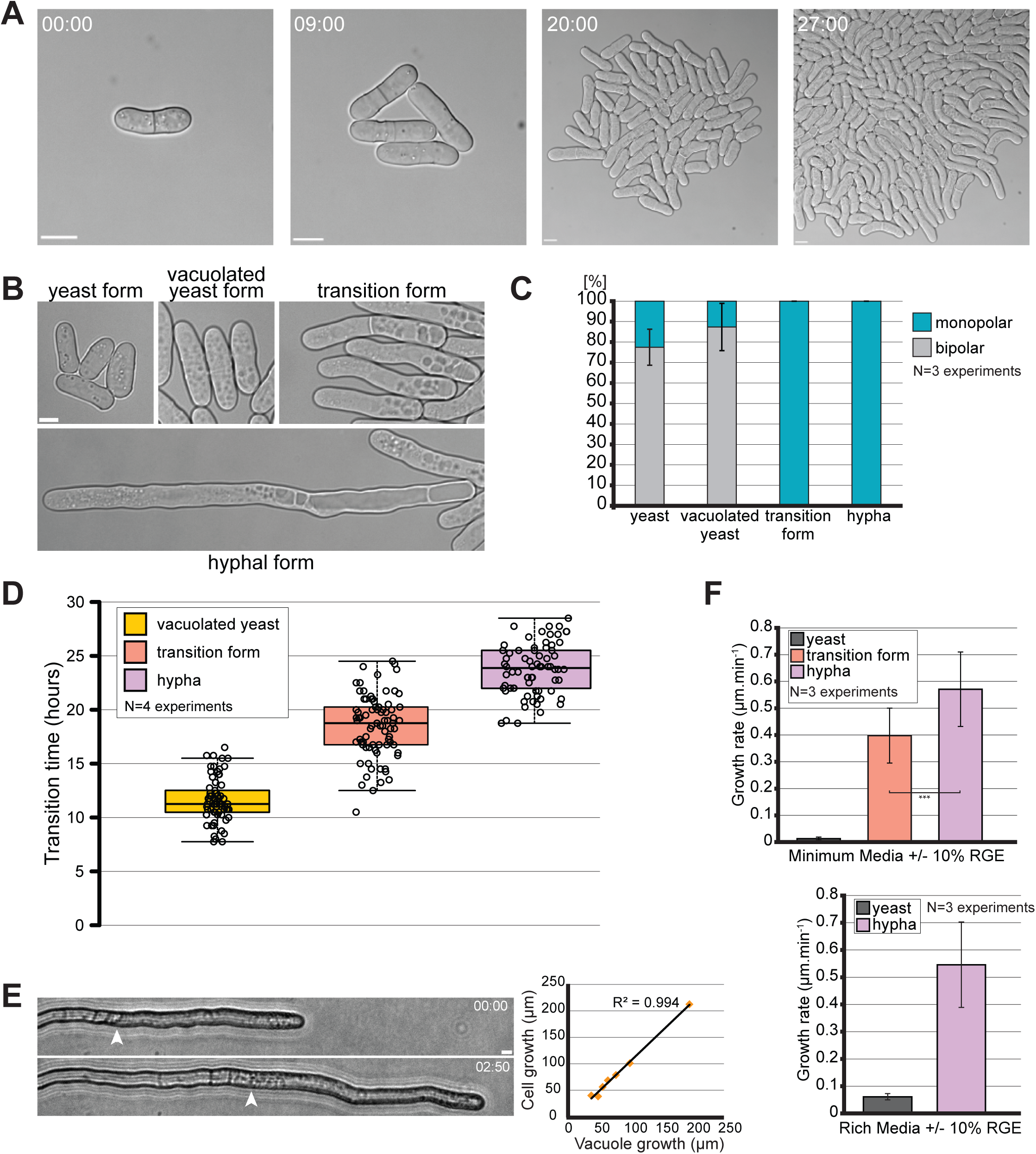
Kinetics of the morphological yeast-to-hypha transition. **A.** DIC microscopy images of mini-colony formation in a microfluidics chamber. **B.** DIC microscopy images of the four identified morphological states of *S. japonicus* during the yeast-to-hypha transition in a microfluidic chamber. **C.** Quantification of monopolar and bipolar cells in the four morphological state of *S. japonicus* (n=51 hyphae and >100 cells for the other states). **D.** Quantification showing time at which each morphological states first appeared after induction with RGE in microfluidic chambers (n>70 cells per state). Box plot shows first and third quartile and median, whiskers extend 1.5 times the interquartile range from the first and third quartile. **E.** Brightfield microscopy images showing a growing hypha on solid media (edge of vacuole shown with arrowhead) and quantification showing a correlation between the growth of the vacuole over time and the growth of an entire hypha over the same amount of time. N=7 hyphae over 5 separate experiments. R^2^= 0.994, linear regression. **F.** Quantification of the growth rate of different morphological forms on minimum and rich media (in minimal medium, n=60 yeasts and >30 cells for the other states; in rich medium, n=60 yeasts and 17 hyphae). Error bars show standard deviations. Time in h:min. Scale bars: 5µm.

### *S. japonicus* does not assemble a classical Spitzenkörper

In an effort to compare *S. japonicus* hyphae to other filamentous fungi and dimorphic yeasts we looked for the presence of a Spitzenkörper at the hyphal tip. The accumulation of vesicles at the Spitzenkörper can be visualized in phase contrast microscopy as a dense spherical organelle (Riquelme and Sanchez-Leon, 2014) or fluorescently labeled with amphiphilic dyes like FM4-64 (Fischer-Parton et al., 2000) or with tagged Rab11 GTPase (Ypt3 in fission yeast), which decorates the vesicles (Cheng et al., 2002). In *S. japonicus*, neither phase contrast imaging, nor FM4-64 showed a spherical signal at hyphal tips. Similarly, GFP-Ypt3 did not reveal a spherical fluorescent signal, though it accumulated at the cortex of hyphal tips, consistent with local vesicle delivery at the site of growth (Fig. 3A). We further tagged other components of the polarization machinery: the exocyst component Exo70 and polarity proteins Bud6 and Spa2, thought to associate with formins, decorated the hyphal tip cortex, but did not form a Spitzenkörper-like structure; the microtubule-delivered Tea1 protein also assumed a localization similar to that described in the cousin species *S. pombe* and did not form a Spitzenkörper-like accumulation (Fig. 3B; movie S2) (Riquelme and Martinez-Nunez, 2016; Takeshita et al., 2008). We conclude that *S. japonicus* does not assemble a classical Spitzenkörper like other filamentous fungi. Moreover, the localization of these polarity factors was similar in the hyphal and the yeast form, with the exception of Bud6, which decorated a notably wider region around the hyphal tip, and comparable to that of their homologues in *S. pombe* (Fig. 3C). This suggests that the transition from yeast to hyphal form occurs without major re-organization of the polarity and trafficking machineries.

**Figure 3.**
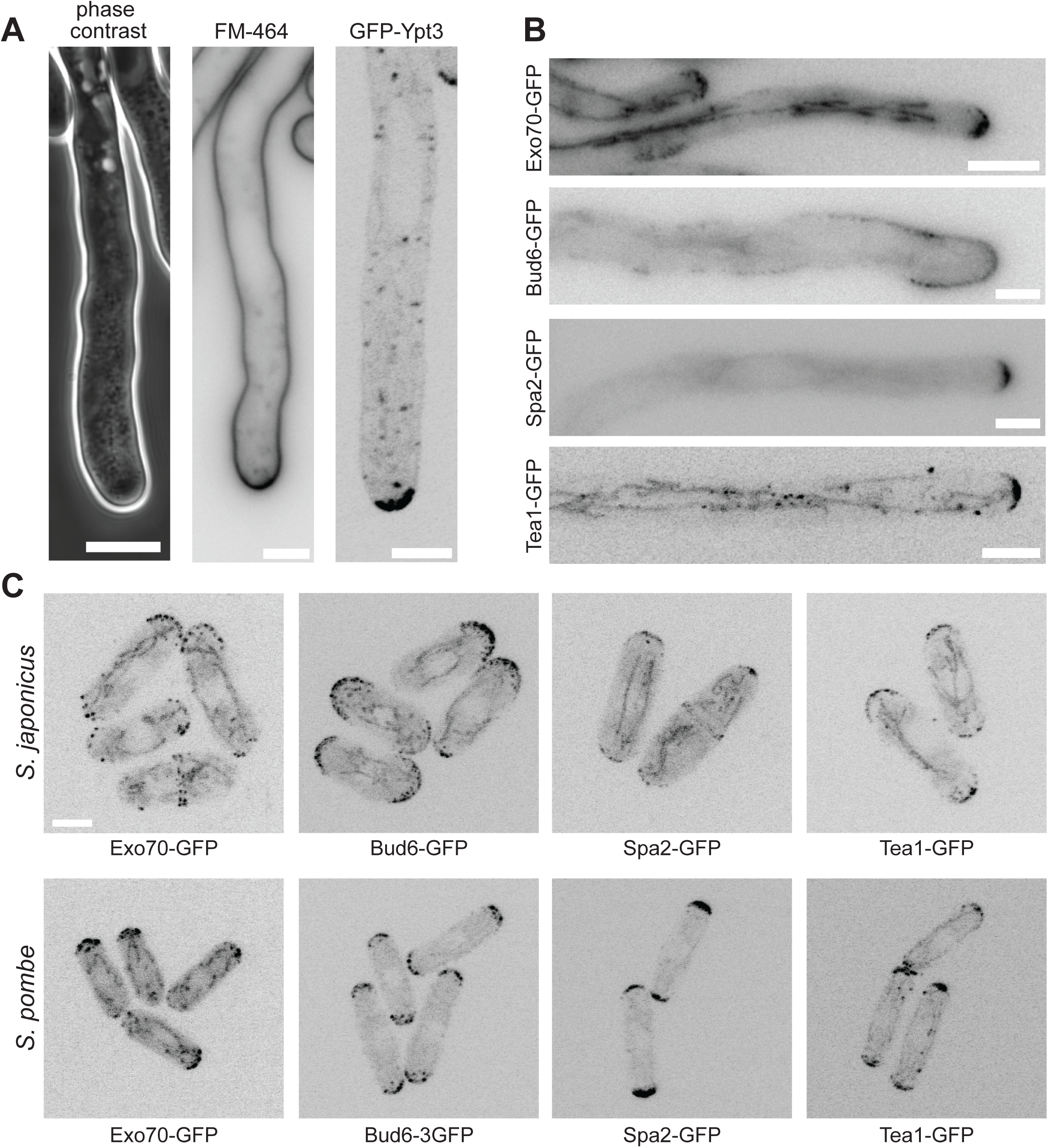
Localization of polarity factors in fission yeasts. **A.** Hyphal tips of *S. japonicus* visualized with a phase contrast objective (left), stained with the amphiphilic dye FM4-64 (middle) and marked with GFP-tagged Rab11 GTPase Ypt3 (right). **B.** Fluorescence images of GFP-tagged polarity proteins Exo70, Bud6, Spa2 and Tea1 at *S. japonicus* hyphal tips. **C.** Fluorescence images of the same polarity proteins in *S. pombe* and *S. japonicus* yeast form. Scale bars: 5µm.

### Actin based trafficking is increased in the hyphal form and is essential for the transition

Although *S. japonicus* does not assemble a Spitzenkörper, live imaging of vesicles tagged with GFP-Ypt3 revealed an important change during the transition from yeast to hypha. Ypt3 vesicles accumulate at the growing tips in both the yeast and the hyphal form of *S. japonicus* and their movement can also be tracked in the cytosol (Fig. 4A; Movie S3-4). Ypt3 fluorescence intensity was significantly increased at hyphal tips compared to yeast cell tips (Fig 4B), suggesting a stronger accumulation of vesicles. This is likely to reflect an increase in membrane traffic to sustain the increase in cell growth. Interestingly, we found that the speed of individual vesicles was also on average significantly faster in hyphae than in yeast (Fig. 4C). Similar fast vesicle speeds and accumulation at cell tip were also observed in the transition form, suggesting that the change in the rate of trafficking happens at the beginning of the transition.

**Figure 4.**
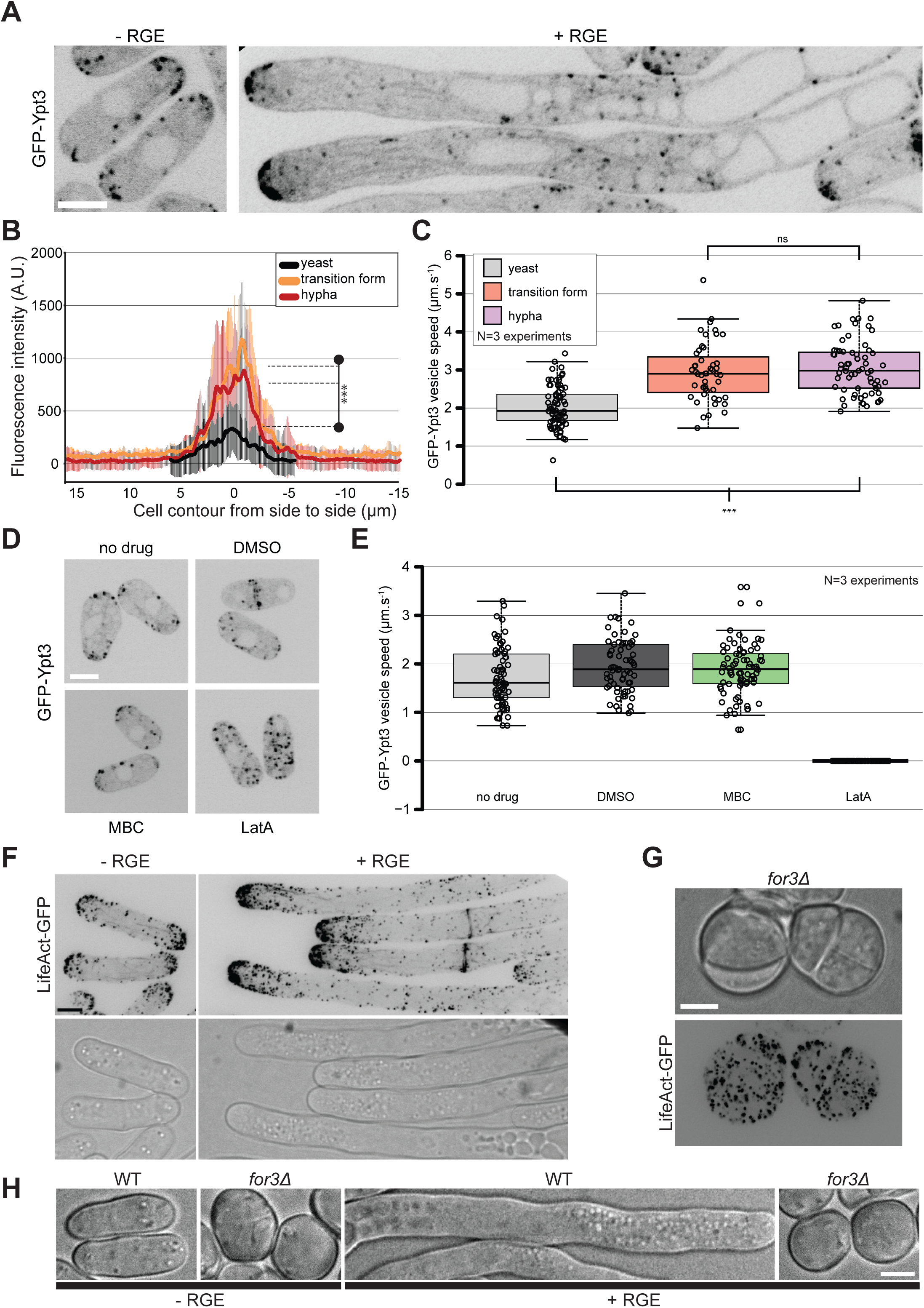
Actin based trafficking is increased in the induced forms. **A.** Middle plane fluorescence images of *S. japonicus* cells expressing GFP-Ypt3 in both non-inducing and inducing conditions. **B.** Quantification of GFP-Ypt3 fluorescence intensity at the tips of yeasts (n=32), transition forms (n=13) and hyphae (n=18). Shaded areas correspond to standard deviations. *** indicates P < 1.2×10^−05^; t. test. **C.** Quantification of vesicle trafficking speed in yeasts, transition form and hyphae. *** indicates P < 4.82×10^−10^; ns indicates P=0.44, t. test **D.** Middle plane fluorescence images of GFP-Ypt3 in cells treated or not with solvent dimethyl sulfoxyde (DMSO), microtubule-depolymerizing MBC or actin-depolymerizing LatA. **E.** Quantification of vesicle trafficking speed in cells treated as in (D). **F.** F-actin localization in *S. japonicus* observed with marker LifeAct-GFP in both yeast and hyphal forms. Images are maximum intensity projections of 16 z-stacks (0.5µm). **G.** Fluorescence images of LifeAct-GFP in *for3Δ* mutants showing absence of actin cables and disorganized patches. **H.** DIC images of wild type and *for3Δ* mutants under non-inducing and inducing conditions. Box plots show first and third quartile and median, whiskers extend 1.5 times the interquartile range from the first and third quartile. Scale bars: 5µm.

Ypt3 trafficking occurred on F-actin, as actin depolymerization with LatA abolished all vesicle trafficking and cell tip localization (Cheng et al., 2002) (Fig. 4D-E; Fig. S1A). By contrast, microtubule depolymerization with MBC had no effect on Ypt3 trafficking. F-actin labeled with LifeAct-GFP was organized in actin patches, cables and rings in *S. japonicus* (Alfa and Hyams, 1990). We noticed an accumulation of actin structures at the tips of growing hyphae coinciding with the increased growth rate for the hyphae (Fig. 4F, see Fig. 2F). Deletion of For3, the formin responsible for actin cable assembly in *S. pombe* (Feierbach and Chang, 2001), led to loss of actin cables in *S. japonicus*, as in *S. pombe*. However, the resulting mutant cells were sicker than their *S. pombe* counterparts (Bendezu and Martin, 2011; Feierbach and Chang, 2001), with impairment in growth and high cell mortality (Fig. 4G, Fig. S1B). *for3Δ* mutant cells did not polarize growth, even in presence of the inducer (Fig. 4H). Thus, *S. japonicus* yeast and hyphal growth rely on transport of vesicles on actin cables for polarized growth, with increased rates of vesicular transport in the hyphal form.

### Microtubules are dispensable for polarized growth of *S*. *japonicus*

We used GFP-Atb2 (alpha-tubulin) to examine the microtubule cytoskeleton. Microtubules form bundles aligned along the length of the cell of both yeast and hyphal forms (Alfa and Hyams, 1990; Sipiczki et al., 1998a). In the yeast form, microtubule organization resembled that described in *S. pombe*, growing from cell middle towards cell ends, sliding along cell sides and shrinking upon touching the cell tip (Fig 5A; Movie S5). In the hyphal form, microtubule bundles were significantly longer, extending over the length of the cytoplasmic segment and were often observed to bend (Fig. 5A; Movie S6). The bundles extended to the hyphal growing tip where they occasionally touched the membrane to deposit polarity factors (see Movie S2). Microtubules also extended through the vacuole-occupying cell segment, though rarely reached the other cell end (Fig 5B; see Fig. 7E). However, short-term microtubule depolymerization with MBC did not impair polarized growth in either yeast (data not shown) or hyphal form (Fig. 5C).

**Figure 5.**
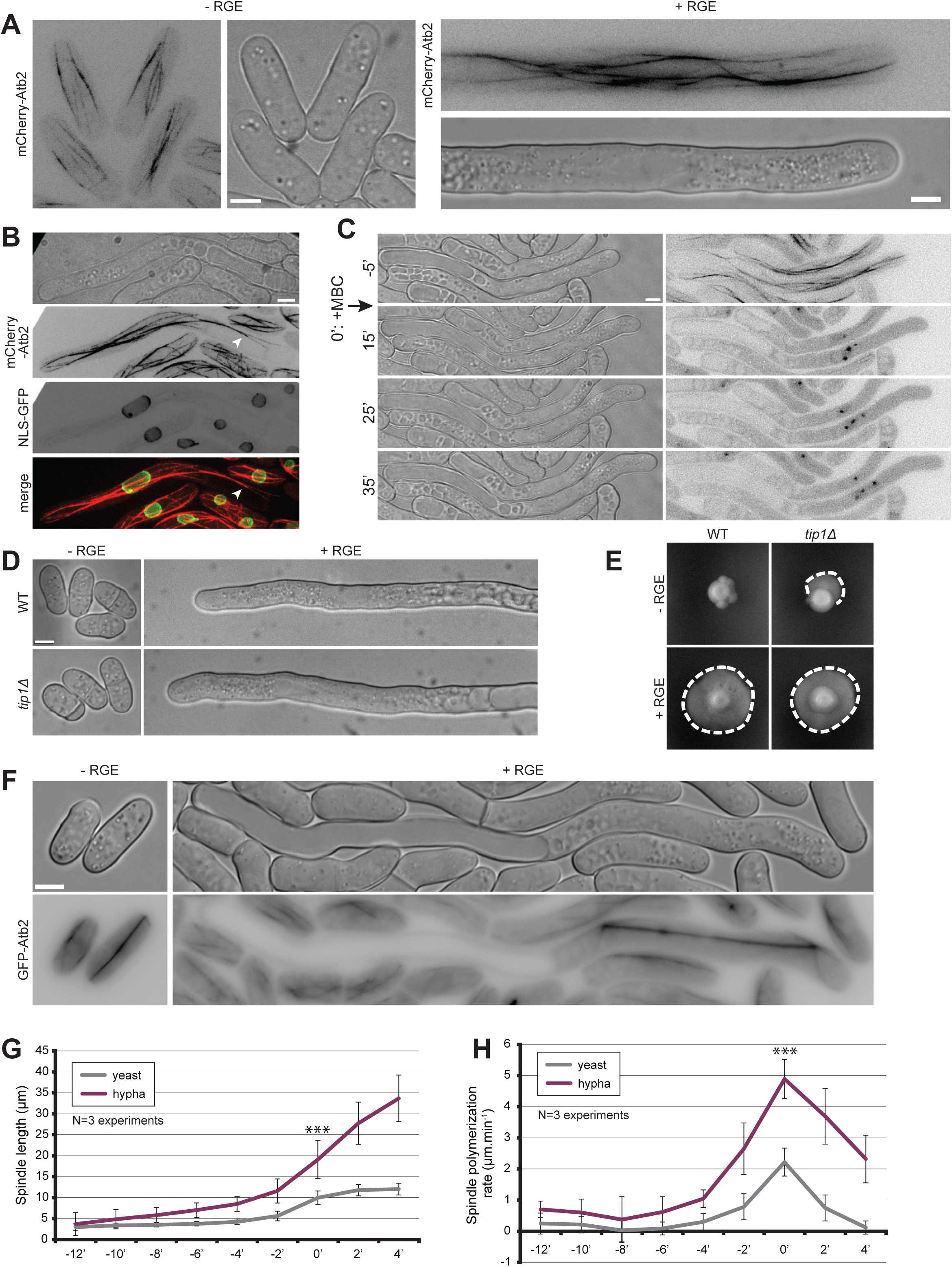
Microtubules are not involved in hyphal growth. **A.** Microtubule organization in *S. japonicus* yeast and hyphal forms expressing mCherry-Atb2. Images are maximum intensity projections of 8-14 z-stacks. **B.** Images of an induced strain tagged with mCherry-Atb2 and NLS-GFP showing microtubules can penetrate the space between the plasma membrane and the vacuole (arrowhead). **C.** Microtubule depolymerization does not perturb hyphal growth. MBC was added at time 0 in a microfluidic chamber. **D.** Wildtype and *tip1Δ* cells grown in microfluidic chambers in inducing and non-inducing conditions. **E.** Wildtype and *tip1Δ* strains grown on solid media in non-inducing and inducing conditions. Dotted lines highlight penetrative filamentous growth. **F.** Mitotic spindles labeled with GFP-Atb2 in yeast and hypha. **G.** Quantification of spindle length over time, aligned on the steepest slope and averaged (n=30 cells per cell type). **H.** Quantification of spindle elongation rates over time. Individual profiles were aligned on the highest rate and averaged (n=30 cells per cell type). *** indicates P < 1.59×10^−10^; t. test. Error bars show standard deviations. Scale bars: 5µm.

Because long-term MBC treatment during the yeast-to-hypha transition gave inconclusive results, likely due to effects on cell proliferation through disruption of the mitotic spindle, we assessed the role of microtubules during the transition by deleting the microtubule plus-tip-associated CLIP-170 homologue Tip1. Though *tip1Δ* cells had some defects, notably in septum positioning, they retained the ability to polarize in the yeast form and to form hyphae within the same timeframe as wild-type (Fig. 5D-E). Hyphae could also still penetrate the agar on solid medium (data not shown). We conclude that microtubules and microtubule plus-tip factors are not important for yeast-to-hypha transition or for hyphal growth.

Microtubules labeling also allowed us to visualize mitotic spindles, which elongated to significantly longer sizes in hyphae than yeast cells: they reach over 30µm in length, almost covering the entire cytosolic hyphal segment (Fig. 5F-G). Interestingly, the rate of spindle elongation was also significantly increased (about 2.5-fold) (Fig. 5H), such that the total duration of mitosis tended to be even shorter in hyphae. We observed also less spindle buckling in hyphae (Yam et al., 2011). This suggests that the rates of microtubule-dependent motors and thus dependent forces, like those of actin-dependent motors driving vesicle movements, are increased in hyphae.

### *S. japonicus* hyphae display complete cell divisions and altered growth controls

While mitosis is common to many filamentous fungi, a more surprising observation is that mitotic divisions were always followed by formation of septa that fully constricted, giving rise to two daughter cells throughout the yeast-to-hypha transition (Fig. 6A; Fig. S2B). This was the case in cells lacking blue-light receptors as well as wild-type cells grown in the dark. Indeed, most filamentous fungi are multinucleated, with some forming septa that do not constrict but help compartmentalize an increasingly complex filamentous network (Mourino-Perez and Riquelme, 2013). Consistent with the completion of cytokinesis, *S. japonicus* remained mononuclear even in the filamentous form (Sipiczki et al., 1998a) (Fig. 6B-C). The nuclei were elongated in the hyphal form with nuclear length correlating well with cytoplasm length, respecting the rule of constant nuclear to cytoplasm ratio (Neumann and Nurse, 2007) (Fig. 6D). This observation, together with the absence of Spitzenkörper described above, sets *S. japonicus* hyphae apart from other filamentous fungi, casting them as more similar to the yeast form than expected.

**Figure 6.**
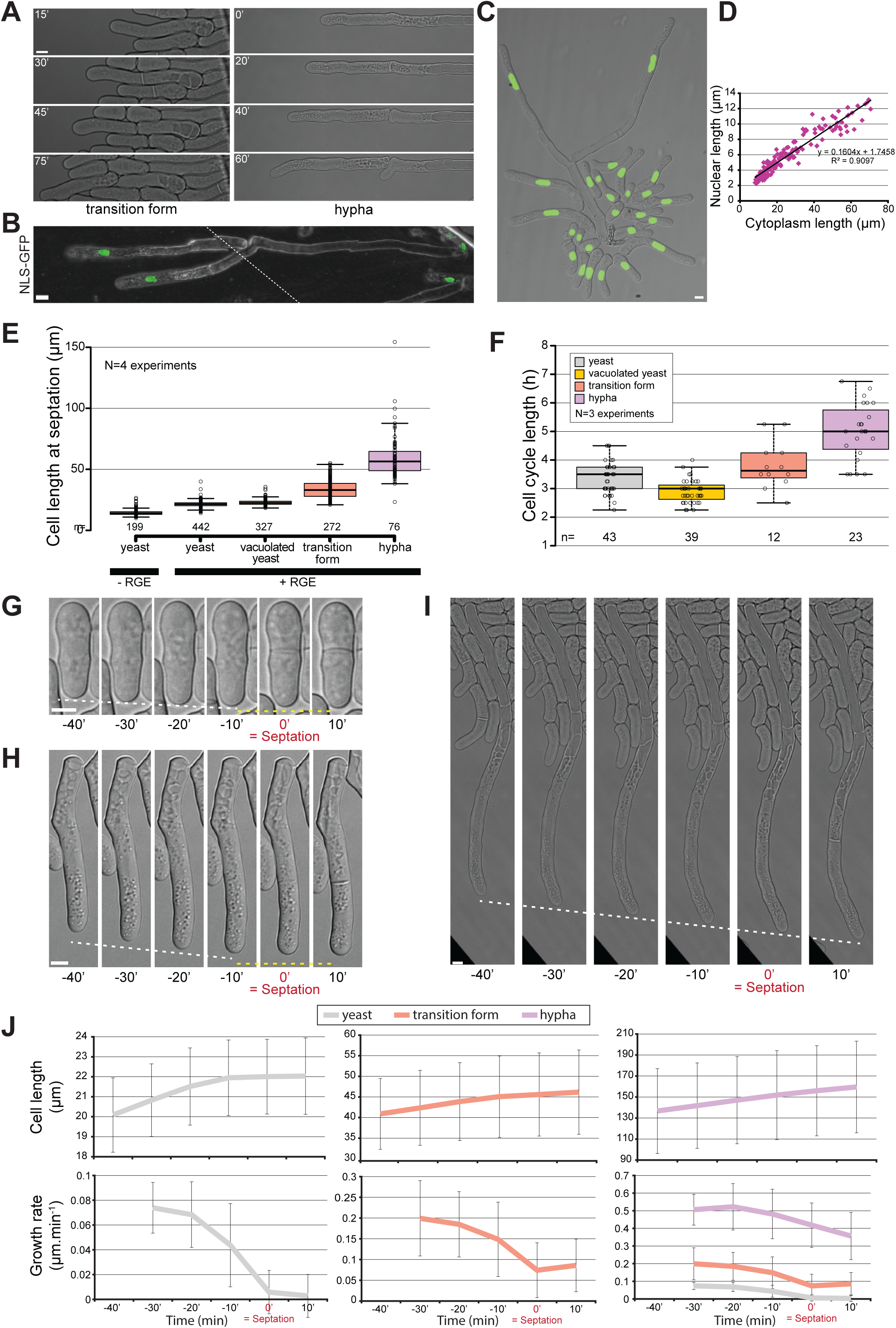
*S. japonicus* produces mononuclear hypha. **A.** DIC images of transition and hyphal forms showing completion of septation. **B.** Tiled confocal microscopy image of hyphae expressing NLS-GFP growing on gelatin plates. Dotted line shows the location of the tiling. **C.** Strain expressing NLS-GFP growing in inducing conditions in a microfluidics plate. **D.** Correlation between nuclear length and cytoplasm length (n=178 cells). **E.** Quantification of cell length at septation in the different morphological forms of *S. japonicus*. **F.** Quantification of cell cycle length in the different morphological forms of *S. japonicus*. **G-I.** DIC images of cell growth in microfluidic plates up to the septation event. White dotted lines show tip growth. Yellow dotted lines show absence of, or reduced, growth. **J.** Analysis of cell length and growth rate over time aligned on septation time for yeast (grey), transition (orange) and hyphal (purple) forms (n> 18 cells per cell type). Box plots show first and third quartile and median, whiskers extend 1.5 times the interquartile range from the first and third quartile. In (J), error bars show standard deviations. Scale bars: 5µm.

Interestingly, in comparison to the yeast form, *S. japonicus* hyphae appeared to show distinct growth control. First, measurement of cell size at division showed a steady increase throughout the yeast-to-hypha transition whereas cell width remained roughly constant, suggesting an alteration in cell size regulation during the transition (Fig. 6E; Fig. S2A). This increase in size was not only due to the fast growth of the transition and hyphal forms as the length of the cell cycle also increased (Fig. 6F). Second, while *S. japonicus* yeast form and *S. pombe* stop growing during septation (Mitchison and Nurse, 1985), we found that the transition and hyphal forms continued to grow (Fig. 6G-I), similar to what is observed in other filamentous fungi (Riquelme et al., 2003). However, in these forms the growth rate decreased during septation, interestingly by a similar absolute value as in yeasts (Fig. 6J). We envisage competition for polarity factors between the growing end and the septation site as a reason for this decrease.

### Highly asymmetric cell division of a fission yeast in *S*. *japonicus*

One fascinating aspect of hyphal division is that this cell division is inherently highly asymmetric (Sipiczki et al., 1998a). Indeed, hyphae (and transition forms) have polarized vacuoles to the back end of the cell and grow in a monopolar manner. Septation always occurred within the cytoplasm-containing cell segment. During cell division, one daughter cell retained the previously built vacuole and little cytosol and paused before growing a branch from the septation point. The other cell inherited most of the cytosol and the hyphal tip, which kept growing as described above. This cell rapidly rebuilt its vacuole close to the septation point (Fig. 7A; Movies S7-S8). Similar behaviors were observed in the transition form (Figure S3). This raises the question of how hyphae position their division site.

**Figure 7.**
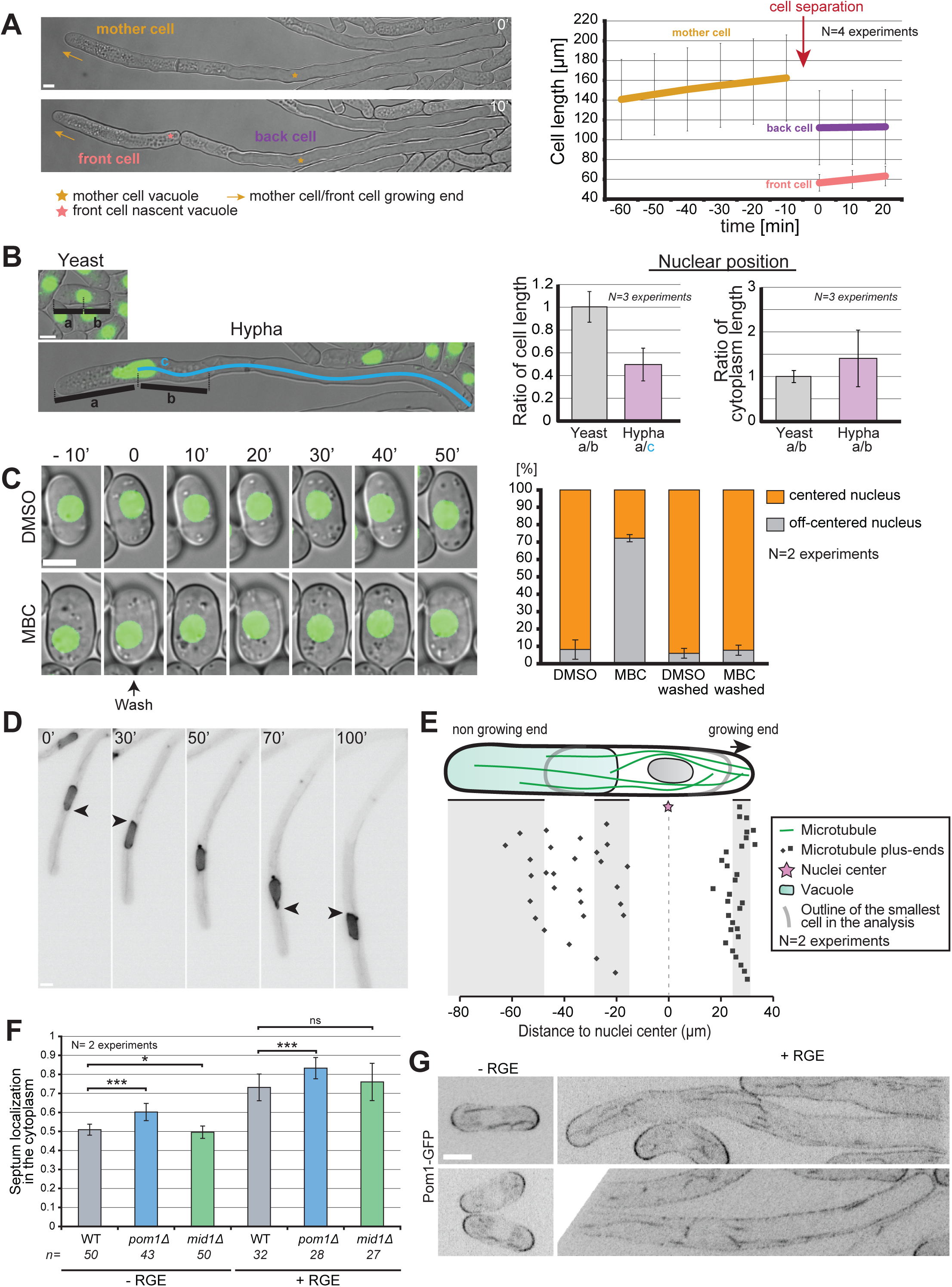
Asymmetric cell division in *S*. *japonicus*. **A**. Hyphae divide asymmetrically, giving rise to a front cell that retains the growing end but has to rebuild a vacuole, and a back cell that retains the vacuole but has to rebuild a growing end (left). Hyphal cell length recording over time aligned on cell separation (right) (n=20 hyphae). **B.** Quantification of nuclear positioning in the cell and in the cytoplasm. Positioning was calculated through ratios as explained in the left panel (n>50 cells per cell type). **C.** Microtubules contribute to nuclear positioning. Cells were grown in microfluidics chambers for three hours with DMSO or MBC and then washed for 50 minutes with EMM-ALU. Nuclear position was quantified before and after the wash in >65 cells per condition. **D.** Fluorescence images of a hypha expressing NLS-GFP showing an example of nuclear shape alteration over time. Arrowheads point to nuclear envelope protrusion indicative of exerted forces. **E.** Schematic of the localization of microtubule plus-ends in the hyphal form. Each dot represents a microtubule plus end position. The nuclear position was used as reference point in all measurements. Shaded areas show the range of positions of the cell front, vacuole front and cell back (n=50 microtubule tips in 10 hyphae). **F**. Quantification of septum position in the cytoplasm of yeast and hyphae, in WT, *pom1Δ* and *mid1Δ* strains. ns: P=0.22; *: P=0.03; ***: P<9.08×10^−08^; t. test. **G**. Middle plane fluorescence images of Pom1-GFP in inducing and non-inducing conditions. In the hyphal form we show both the front and the back of cells. Scale bars: 5µm.

We first investigated the mode of nuclear positioning. In *S. pombe*, the nucleus is positioned at mid-cell due to microtubules anchored at the nuclear envelope exerting pushing forces against both cell poles (Daga et al., 2006; Tran et al., 2001). In *S. japonicus* yeast cells, nuclei were at mid-cell, as in *S. pombe*. By contrast, in hyphae, nuclei were not at mid-cell, but were displaced towards the cell front because the vacuole occupies the back of the cell. However, they were also not centered within the cytosolic segment, but displaced towards the vacuole (Fig. 7B). To examine the role of microtubules in nuclear positioning, we performed depolymerization experiments. In the yeast form, after over three hours of depolymerization with MBC about 70% of the cells showed a misplaced nucleus. 50 min after washout, most nuclei centered again to the cell middle, indicating microtubules control nuclear positioning (Fig. 7C). Similar experiment in the hyphal form proved to be challenging, but examination of nuclear morphology during hyphal growth showed frequent nuclear shape deformation indicative of forces exerted on the nuclear membrane (Fig. 7D). Nuclear envelope deformations were seen on both sides of the nucleus, suggesting that microtubules exert forces from both sides. We noted above that microtubules penetrate the vacuole-occupied cell segment (see Fig. 5B). The quantification of microtubule plus-end positioning within the cell showed a strong accumulation close to the hyphal tip, where accordingly to data in *S. pombe* they are expected to exert pushing forces. On the vacuole side, most microtubules were able to penetrate the space between the vacuole and the plasma membrane, though the majority ended within the first half of the vacuole length, suggesting that the pushing force may be partly dissipated (Fig. 7E). Thus, we hypothesize that the force exerted on the nuclei by microtubules growing towards the vacuole is weaker than that produced by microtubules growing towards the hyphal tip, leading to the observed bias in nuclear positioning towards the vacuole.

Similar to nuclei, hyphal septa were always positioned within the cytosolic cell segment, though they were off-centered towards the vacuole (Fig. 7F). This position was unaltered in *mid1Δ* hyphae, indicating that, as in the yeast form (Gu et al., 2015), Mid1 is not involved in septum positioning in *S. japonicus*. However, the pre-divisional nuclear position did not predict the septum position, which was better, though not perfectly predicted by the middle of the anaphase spindle (Fig. S4). We note that, as in the yeast form, hyphal mitosis is semi-open (Fig. S4) (Yam et al., 2011). These observations suggest that positive signals for septum assembly may be conferred by the spindle.

Pom1 kinase constrains septum placement to mid-cell in *S. japonicus* yeast form, as it does in *S. pombe* (Gu et al., 2015). Similarly, we found that *pom1Δ* hyphae showed septum mispositioning, where the septum was excessively displaced towards the vacuolar segment (Fig. 7F). In *S. pombe*, this has been attributed to Pom1 gradients from cell poles exerting negative control to prevent septation at cell tips. Similar distribution is apparent in *S. japonicus* yeast cells (Fig. 7G). Curiously, in hyphae, though Pom1-GFP accumulated at cell poles, it was also very distinctly present along cell sides, and did not form an obvious long-range concentration gradient (Fig. 7G). This raises the question of how Pom1 conveys positional information for septum placement.

## Discussion

### A mycelium formed of single cells

It is believed that all Ascomycetes descend from a common filamentous ancestor (Berbee and Taylor, 1993). The fission yeasts form an early-diverging ascomycete clade, amongst which *S. japonicus* is the most divergent, suggesting that *S. japonicus* has retained an ancestral ability to filament present in the last common fission yeast ancestor (Sipiczki, 2000). In the fungal kingdom, filamentation leads to the formation of a mycelium, a complex multicellular network that underlies fungal spread and can reach several meters across (Islam et al., 2017; Smith et al., 1992). Mycelia are typically formed of a single, large common cytosol, which completely lacks septa in lower fungi, or is compartmentalized by incomplete, pore-containing septa in higher fungi (Steinberg et al., 2017). As a result, mycelia can typically be considered as a multinucleated syncytia. Hyphal fusion, or anastomosis, further increases the level of interconnectedness in fungi (Heaton et al., 2012; Read et al., 2010; Read et al., 2009), and this plays an important role in nutrient exchange within the fungus (Simonin et al., 2012). Although having a single cytoplasm extended over such lengths can appear risky, septate hyphae can easily seal off their septa, an absolutely vital process to prevent loss of cytoplasm in case of damage on the mycelium (Riquelme et al., 2018), in case of unfavorable environment (van Peer et al., 2010), or during aging (Bleichrodt et al., 2015). In this work we offer a description of a different kind of mycelium in *S. japonicus*.

The dimorphism observed in *S. japonicus* leads to the formation of extremely polarized and elongated single cells that are highly invasive of solid substrate similar to what is displayed by other filamentous organisms. In this work, we describe fruit extracts as novel inducers for the yeast-to-hypha transition in *S. japonicus*. Previously, nutrient starvation and DNA damage had been described as methods of induction of dimorphism in this fungus. Our method of induction appears to be specific for *S. japonicus*, as it did not trigger morphological transition in other fission yeasts or in *S. cerevisiae*, which forms penetrative pseudohyphae in response to nitrogen starvation (Gimeno et al., 1992). We note that filamentous forms have been reported for *S. pombe* (Amoah-Buahin et al., 2005), though RGE did not promote their formation. Cues triggering dimorphism are very varied in fungi; for example *Candida albicans* undergoes hyphal formation through a multitude of signals including serum and pH (reviewed in (Sudbery, 2011)), and *Ustilago maydis* filamentation can be triggered by sexual pheromones and the resulting dikaryotic hyphae will infect maize plants (Nadal et al., 2008). Interestingly, *S. japonicus* was isolated from both strawberries and grape extracts (Wickerham and Duprat, 1945; Yukawa and Maki, 1931), both of which we have shown to promote hyphal growth. This raises the question of its natural habitat and which morphological form it adopts in the wild. The hyphae produced by *S. japonicus* grow in average at 0.58µm.min^−1^, a rate similar to what is observed in true filamentous fungi *Aspergillus nidulans* (0.5 µm.min^−1^, (Horio and Oakley, 2005)) and fellow dimorphic yeast *C. albicans* (0.75 µm.min^−1^ (Gow and Gooday, 1982)). Thus, in appearance, the transition of *S. japonicus* leads to the formation of a macroscopic mycelium.

However, in contrast to other mycelia, our study reveals that the *S. japonicus* mycelium is fragmented. Indeed, *S. japonicus* hyphae not only place septa, similar to what is observed in septate hyphae of higher fungi, but also fully divide after mitosis. As a result, all hyphae are mononuclear, a very unusual feature for a filamentous organism. After hyphal division, both front and back cells resume growth, with the back one resuming growth at an angle behind the recently formed septum, which superficially resembles a branch point. However, we never observed true branching. The lack of a Spitzenkörper in *S. japonicus* hyphae is another important point of divergence from other filamentous Ascomycetes. Indeed, the Spitzenkörper, a vesicle supply center that promotes and orients hyphal growth, is largely associated with Ascomycetes and Basidiomycetes septate hyphae, but is usually absent from early-diverging fungal lineages. Instead, we find that *S. japonicus* hyphae accumulate secretory vesicles at the growing tip in a less clustered pattern, similar to what was observed in yeast growth and filamentous Zygomycetes species (Grove and Bracker, 1970; McClure et al., 1968; Roberson et al., 2010). Finally, the strict dependence of hyphae on actin-based transport and independent from microtubules also cast it apart from most other filamentous fungi, which use microtubules for long-range transport (Egan et al., 2012). These characteristics raise the question of whether the *S. japonicus* filamentous form should be considered true hyphae or pseudohyphae. Its complete septation and mononuclearity and its lack of Spitzenkörper are pseudohyphae characteristics. However, the very large vacuoles in *S. japonicus* filaments are a feature of hyphae. Thus, the novel filamentation process of *S. japonicus* described here, which does not rely on the typical landmarks, represents an intermediate form of filamentation and participates to the wide variety of filamentous forms in fungi.

### Asymmetrical division in fission yeasts

An interesting aspect of the yeast-to-hypha transition in *S. japonicus* is the conversion of a symmetrical to an asymmetrical system. In the yeast form, the cell grows at both poles and divides in the middle, generating two apparently equivalent daughters, similar to the case of *S. pombe*. By contrast, *S. japonicus* hyphae are morphologically and functionally very asymmetrical. Division yields a front cell that contains most of the cytoplasm and the unique growing tip, and is shorter than the back cell, which is largely filled with an ever-growing vacuole and has to re-initiate growth with a delay. This asymmetrical conversion is already apparent early in the transition when cells switch to a monopolar mode of growth, which coincides with the accumulation of initially fragmented vacuoles to the back of the cell. Because *S. japonicus* can be easily induced to switch from a symmetrical to an asymmetrical division, the signals and mechanism of this conversion are rich grounds for future investigations.

One aspect we explored in a little more detail is the question of nuclear and septum positioning. While nucleus and septum are placed at mid-cell in the yeast form, in wild type hyphae we have shown that both are displaced away from the middle. The positioning mechanism for the nucleus can be inferred from work in *S. pombe*, which showed that microtubules anchored at the nuclear membrane exert pushing forces against cell poles, such that force balance is achieved when the nucleus is centered in the cell (Daga et al., 2006; Tran et al., 2001). Microtubules also exert forces for nuclear positioning in *S. japonicus*, as illustrated by the observations that decentered nuclei are re-centered upon microtubule regrowth in yeast and the nuclear envelope is deformed by microtubules in hyphae. However, the nucleus is positioned neither in the middle of the hypha nor in the middle of the cytoplasmic region, but closer to the vacuole. We suggest that microtubule-dependent pushing forces position the nucleus, as in *S. pombe*, but that forces only become balanced at a non-medial position due to force dispersion when microtubules encounter the less rigid vacuole at the back of the cell compared to the rigid cell wall at the front.

The positioning of the septum is more mysterious. In *S. pombe*, septum positioning at mid-cell is widely thought to rely on two complementary signals – a positive nuclear signal transmitted through the anillin-like protein Mid1, and a negative cell pole signal dependent on Pom1 kinase (Celton-Morizur et al., 2006; Chang et al., 1997; Huang et al., 2007; Padte et al., 2006; Sohrmann et al., 1996). In *S. japonicus* yeast cells, recent work showed that Mid1 plays no significant role in septum positioning (Gu et al., 2015). We confirm this in hyphae, as septum position is not altered in *mid1Δ* cells. Consistently, we find that the septum position is poorly predicted by the pre-divisional nucleus, indicating that septum position is defined at later time than in *S. pombe*. The septum was closer to but not perfectly predicted by the center of the anaphase spindle, suggesting positioning signals may be more similar to those used in metazoan cells, where the spindle is the key determinant (Oliferenko et al., 2009). By contrast, Pom1 kinase regulates division site positioning in both yeast and hyphae ((Gu et al., 2015) and this work). For this function, Pom1 was proposed to form an inhibitory concentration gradient from the cell poles that counteracts the localization of medial cytokinetic node precursors, which in consequence preferentially form at mid-cell (Bhatia et al., 2014; Celton-Morizur et al., 2006; Padte et al., 2006; Rincon et al., 2014). The long distance between the growing pole and the septum (over 50µm on average) calls this view into question, at least in hyphae. Indeed, although Pom1-GFP distribution in yeast cells was very similar to that described for *S. pombe* (Hachet et al., 2011), in hyphae it was only mildly enriched at tips and decorated most of the plasma membrane without forming an obvious long-range concentration gradient from the tip to the site of division. The cortex at the back of the cell, occupied by the vacuole, was also strongly decorated by Pom1, without enrichment at the back cell pole. This raises the question of how Pom1 conveys positional information for division in hyphae.

### Size control

The *S. japonicus* yeast-to-hypha transition also provides an excellent system to study principles of size control. As the cell dramatically lengthens during the transition, several aspects of growth and division control are notably altered. *S. pombe* is arguably one of the best-studied systems for size control, due in part to its highly reproducible length at division. Measurements of cell size homeostasis concur in proposing that the system functions as a sizer (Wood and Nurse, 2015), with recent work suggesting that the key dimension informing on division timing is the surface area (Pan et al., 2014). Whether this holds true for *S. japonicus* yeast form is currently not known, but the much longer cell length at division of transition and hyphal cells indicates a profound change or relaxation in the mode of size control. Because cell cycle length is also increased during the transition, a simple timer model, where increased cell size would be acquired due to faster growth during a set time, is also unsatisfactory. However, we note that the often observed correlation between cell and nuclear size (Jorgensen et al., 2007; Neumann and Nurse, 2007; Webster et al., 2009) is also present in *S. japonicus*. This correlation is spectacular, covering over 7-fold variations in size. The observed correlation occurs when nuclear size is compared with the cytoplasmic hyphal compartment rather than the whole cell, whose length varies according to the size of the vacuole. This is in agreement with data in *S. pombe* that support the idea that cytoplasmic volume determines nuclear size (Neumann and Nurse, 2007). These observations suggest that any size control in hyphae may monitor cytosolic and/or nuclear volume excluding vacuoles rather than length or surface area.

Many aspects of cell physiology are faster in the hyphae. First, polarized growth is over ten-fold faster, despite cell width remaining roughly constant. This indicates the surface of the cell tip is not the limiting factor for polarized growth and that growth material must be supplied at an increased rate. Second, we find that secretory vesicles indeed display faster linear movements in hyphae. This increase in transport rate may contribute, but is unlikely to fully explain the increase in growth rate, because it is considerably milder (about 1.5-fold). However, it indicates that myosin (likely myosin V) motors inherently move faster, or are less impeded in their progression in the hypha. Third, it is intriguing that spindle elongation rates are similarly increased (about 2.5-fold). As spindle elongation primarily relies on the action of kinesin motors, this suggests that kinesin motor speed is increased by a similar factor as myosin. As the duration of anaphase (as measured by the spindle elongation phase) is similar in yeast in hyphae, this produces much longer spindles in hyphae. Finally, we found that, in contrast to the yeast form, polar growth does not cease during hyphal division. In *S. pombe*, antagonism between two signaling pathways, the SIN and MOR pathways, is thought to control the alternation between cytokinesis and polarized growth (Ray et al., 2010). This suggests that crosstalk between these two signaling pathways, and more generally between growth and division, is altered upon hyphal transition.

In summary, our detailed description of the yeast-to-hypha transition in *S. japonicus* provides the founding work for addressing important fundamental cell biological questions. The identification of fruit extracts as inducer permits a simple stress-free induction to study an important morphological transition. In particular, the conversion of the cell from a symmetrical to an asymmetrical division system and the massive changes in size in a single cell promise to reveal novel principles in division, growth and size control.

## Materials and Methods

### Strains and media

The original wild-type auxotrophic *S. japonicus* strains were kindly provided by H. Niki (Furuya and Niki, 2009). The *S. japonicus* open reading frames (ORF) used in this study are the following Atb2 (SJAG_02509), For3 (SJAG_04703), Spa2 (SJAG_03625.5), Bud6 (SJAG_04624.5), Tea1 (SJAG_01738), Exo70 (SJAG_04960), Ypt3 (SJAG_03915), Tip1 (SJAG_002695), Pom1 (SJAG_02392), Mid1 (SJAG_01143), Wcs1 (SJAG_02860) and Wcs2 (SJAG_05242). Cells were typically cultured in rich media (YE: yeast extract, 5g; glucose, 30g/liter) for agar plate based experiments and in Edinburgh minimal medium (EMM) supplemented with the appropriate amino acids (EMM-ALU) for microfluidic-based experiments. The fast growth of *S. japonicus* yeast cells in YE would entirely fill the microfluidic plates before transitioning to hyphae in presence of the inducer, which is why we chose to work with minimum media in this case. Red grape extract was obtained from blending 500g of red grapes; the current batch of inducer was made from Crimson seedless grapes from Brazil. The blended grapes were placed in 50ml Falcon tubes and centrifuged at 10000rpm for 25min at room temperature (Eppendorf A-4-62). After recovery of the liquid supernatant by pipetting, the extract was placed in clean 50ml tubes and centrifuged a second time (10000rpm, 15min). Depending on the batch of grapes this step was sometimes repeated. The grape extract was then filtered through a 0.22µm filter (Millipore), aliquoted and kept at −20°C for a maximum of 2 years before degradation of the inducing capabilities. Hyphal formation was induced by adding 10% of red grape extract (RGE) to liquid or solid media. Crosses were done on SPAS media as previously described (Furuya and Niki, 2009) and strains were selected by random spore analysis.

### Strains construction in *S. japonicus*

The genome has been sequenced (Rhind et al., 2011) and is available at: http://fungidb.org/fungidb/

We used homologous recombination to introduce GFP or mCherry fluorescent markers, or to delete a gene. Most of the plasmids constructed in this study were derived of pJK-210 backbone containing the ura4 cassette from *S. japonicus*. This plasmid was constructed and kindly provided by Dr. S. Oliferenko (Crick Institute, London).

To create a gene deletion, the 5’ untranslated region (UTR) was linked with an inverted 3’ UTR fragment (1Kb each at least), separated by a unique restriction site, by PCR stitching and cloned in a ura4+-containing pJK210 plasmid. Homologous recombination in *S. japonicus* is only efficient when homology between the fragment to be integrated and the genomic locus extend to the very end of the fragment. We thus chose 3’ and 5’ UTR fragments in such a way that stitching reconstitutes a blunt restriction enzyme site. Typically, we reconstructed a SmaI restriction site (CCCGGG) by choosing a 5’UTR region to amplify that started with GGG and a 3’UTR region that ended with CCC. Linearization of the plasmid and transformation led to gene replacement by the plasmid through homologous recombination.

To create Ypt3 fluorescently tagged N-terminally with GFP the same procedure was used and the linked 3’ and 5’ UTR regions were inserted before the GFP coding sequence containing no stop codon. The ORF, with a stop codon, was inserted after the GFP and the plasmids were linearized with SmaI reconstructed between the stitched 3’ UTR and 5’ UTR regions.

To create a N-terminally tagged protein mCherry-Atb2, we generated a plasmid containing the putative promoter of Atb2 (we amplified 1.4Kb upstream of the Atb2 ORF) followed by the mCherry coding region without the stop codon and the ORF of Atb2. This plasmid was linearized with AfeI located in the ura4 coding sequence on the plasmid and was transformed in a strain with a mutated *ura* locus where it reconstructed a functional *ura* gene.

To create a protein tagged with a fluorescent marker at the C-terminus, we amplified at least 1kB of the end of the ORF containing a restriction site and no stop codon, and inserted it in the plasmid containing GFP or mCherry coding sequences. After linearization, the plasmid was transformed and inserted in the native loci of the genes of interest.

Transformation was done as previously described (Aoki et al., 2010). Briefly, the cells were grown to exponential phase and then washed in ice-cold water and 1M sorbitol. After incubation with 1M DTT, the cells were put in contact with at least 300ng of linearized plasmid. Transformation of the cells was achieved through electroporation in 0.2cm cuvettes, with those exact settings: 2.3KV, 200Ω, 25µF (Gene Pulser II, Biorad). Cells were left in liquid YE medium overnight to recover and plated the next day on selective media (EMM-AL, lacking uracil).

In the case of the construction of Pom1-GFP strain, we linked together a fragment of the 3’ UTR region with a fragment of the end of the ORF without the stop codon and we cloned the stitched fragment in a pFA6a-GFP-kanMX plasmid in front of the fluorescent marker. After linearization the plasmid was transformed in a wildtype prototroph strain following the same transformation protocol and selected on YE-G418 plates.

All strains were checked for correct insertion of the plasmid with diagnostic PCR and in the case of deletions we also used primers inside the coding region and confirmed the ORF was properly deleted.

#### Microscopy imaging

Wide-field microscopy was performed on a DeltaVision platform (Applied Precision) composed of a customized inverted microscope (IX-71; Olympus), a 60x/1.42 NA oil objective, a camera (CoolSNAP HQ2; Photometrics or PrimeBSI CMOS; Photometrics), and a color combined unit illuminator (Insight SSI 7; Social Science Insights). Figures were acquired using softWoRx v4.1.2 software (Applied Precision). Spinning-disk microscopy was performed using an inverted microscope (DMI4000B; Leica) equipped with an HCX Plan Apochromat 100×/1.46 NA oil objective and an UltraVIEW system (PerkinElmer; including a real-time confocal scanning head [CSU22; Yokagawa Electric Corporation], solid-state laser lines, and an electron-multiplying charge-coupled device camera [C9100; Hamamatsu Photonics]). Stacks of z-series confocal sections were acquired at 0.5-to-1µm intervals using Volocity software (PerkinElmer). Confocal microscopy tile scan images were acquired with a Zeiss laser scanning microscope (LSM 710) mounted with an EC Plan-Neofluar 40X/1.30NA oil objective. Images of growing yeast colonies on agar plates were imaged with a Leica MZ16 FA stereomicroscope (magnification 80-100 times). Images of growing hyphae on agar pads were imaged with a Leica brightfield microscope mounted with a 20X air objective. Phase contrast imaging was acquired on Nikon Eclipse Ti microscope, mounted with a 100X phase contrast objective.

#### Hyphal transition experiments

Cellasic ONIX microfluidics system was routinely used to image the transition (CellAsic system, Millipore, USA, (Lee et al., 2008)). To image the yeast form, cells were grown overnight in 3ml of liquid EMM-ALU to OD600 = 0.4 and then loaded in the plate 2 hours prior to imaging to give them time to settle. To image the hyphal form, the cells were grown in liquid EMM-ALU overnight up to an OD600 of 0.1-0.2, loaded in the microfluidics plate and grown in EMM-ALU-10%RG for 12 to 15hours before imaging at 3 psi (20.7 kPa) in complete darkness (plate surrounded with aluminum foil) to induce hyphal formation. To observe the transition on solid agar plates (agar bacteriological, Oxoid, LP0011), we typically cultured *S. japonicus* overnight in 3ml EMM-ALU and let the cells grow to exponential phase. Cells were then spun down and concentrated before being plated on solid agar plates containing 10% of red grape extract. Hyphal growth was assessed 4 to 12 days later, as indicated. To assess growth rate in rich media, cells were grown on YE-2%agar microscopy pads supplemented or not with RGE for 12 hours before imaging. For the confocal microscopy imaging, *S. japonicus* hyphae were grown on EMM-ALU 12% gelatin (Sigma-Aldrich, #48723) plates supplemented with 10% RGE for 8 days before a piece of gelatin containing hyphae was cut out and mounted on a slide for imaging.

#### Drug treatments and FM4-64 staining

To depolymerize actin, we used a 20mM stock of Latrunculin A (LatA) dissolved in dimethyl sulfoxide (DMSO) to exponentially growing cells to a final concentration of 200µM. Methyl benzimidazole carbamate (MBC, Sigma) was used for the depolymerization of microtubules. A stock solution at 2.5mg/ml in DMSO or ethanol was made freshly on the day of the experiment and exponentially growing cells were treated at a final concentration of 25µg/ml for 10min at 30°C. To depolymerize microtubules in the microfluidics chambers we flowed in EMM-ALU containing 25µg/ml at 3psi, which led to total depolymerization within 10-15 min, similar to the timing observed in liquid cultures. To wash the drug away, we flowed in EMM-ALU at 3 psi and recovery of the cytoskeleton was observed within a few minutes. FM4-64 stainings on growing hyphae were performed in microfluidics chambers as previously described (Fischer-Parton et al., 2000).

#### Treatment of RGE

To identify the molecule in the red grape extract responsible for the morphological transition, we submitted the RGE to a variety of physical and enzymatic treatments. RGE was treated with 16u of proteinase K (NEB, 800u/ml), 20u and 100u of DNAse I (NEB, 2000u/ml) and 1000u of RNAse If (NEB, 50000u/ml). RGE was boiled to 95°C for 20min. RGE was subjected to chloroform/methanol mix (1:1) and left to phase separate overnight at −20°C. Both resulting aqueous and organic phase were dried with a nitrogen stream and subsequently suspended in PBS 1X. All treated RGE were then included in solid agar plates and tested for hyphal inducing capabilities (see Table 1).

#### Supplementation of YE media

To identify the molecule in the red grape extract responsible for the morphological transition, we also tried supplementing rich media with several likely molecular components of RGE and assess for hyphal formation. Liquid YE media was individually supplemented with an additional 40g.L^−1^ glucose, with 100g.L^−1^ fructose, with concentrations ranging from 2mg.L^−1^ to 40mg.L^−1^ resveratrol (Enzo Lifesciences) or with 270mg.L^−1^ ascorbic acid. All supplemented YE media were then included in solid agar plates and tested for hyphal inducing capabilities (see Table 1).

#### Tropism assay

Cells were grown to exponential phase in YE and concentrated 10 times. On large petri dishes (120mmx120mm) containing YE-2%agar, we drew a 6 cm line in the middle of the plate on which 50µl of cells were deposited. 2 cm away from the center of the line, we dropped a small piece of chromatography paper (Whatman, 0.34mm, #3030-917) on which we pipeted 50µl of RGE or YE (control). Plates were covered with aluminum foil and left at 30°C for 12 days before quantification of the area of hyphal growth. A ratio of positive tropism (growth towards the filter) on negative tropism (growth away from the filter) was calculated for both the experiment and the control plates for each plate and averaged over two experiments.

#### Identification of the different morphological forms

In all our experiments we determined the stage of the morphological transition by looking at the polarity stage (monopolar/bipolar), the general localization of the vacuoles (all around the cytoplasm or already polarized at one pole) and the number of vacuoles.

<u>Yeast</u>: bipolar growth, no apparent vacuoles

<u>Vacuolated yeast</u>: bipolar growth, vacuoles all around the cytoplasm

<u>Transition form</u>: monopolar growth, many small vacuoles polarized at the non-growing end

<u>Hypha</u>: monopolar growth, one or two larger vacuole(s) at the non-growing end

#### Bipolar/monopolar quantification

On DIC (differential interference contrast) movies we recorded how many cells had one or two poles growing in the different forms of the morphological transition.

#### Growth rate calculations

On DIC movies we calculated the growth rate by measuring the change in cell or vacuole length over time. Cell growth rates on minimum media were calculated from cells growing in microfluidics plates in inducing and non-inducing conditions and were averaged over three experiments. Cell growth rates on rich media were calculated from cells growing on agar pads in inducing and non-inducing conditions and were averaged over three experiments. The correlation of cell growth and vacuole growth was calculated from cells growing on agar pads in inducing conditions, each point in the graph representing a distinct hypha for which we calculated the extension length of the cell and the vacuole over the total time of the movies. We note that movies were of different length but it clearly demonstrates the correlation between both extension lengths.

#### Fluorescence levels

On spinning disk medial focal planes of GFP-Ypt3 tagged cells (150 ms exposure, 100% laser power) we calculated the fluorescence intensity at the tips of yeasts and transitioning cells by drawing a segmented line of 15 pixels in width around the cell periphery. We subtracted background noise averaged from two different fields of view per experiment. Fluorescence profiles were aligned to the geometric cell tip and averaged by cell type and over three experiments.

#### Quantification of vesicle speed

On spinning disk movies we manually measured the total trajectory of individual Ypt3 dots and derived the rate by dividing by the total time. Data was averaged over three experiments and averaged by cell type. Box plots were generated with http://shiny.chemgrid.org/boxplotr/

#### Quantification of lengths

Spindle length was measured at each timepoint from apparition to complete elongation on epifluorescence movies of cells tagged with GFP-Atb2. Cell length was measured by drawing a line across the cell length from cell pole to cell pole on transmitted light images on septating cells. Box plots were generated with http://shiny.chemgrid.org/boxplotr/. Nuclear length was calculated by drawing a line across the nuclei tagged with NLS-GFP construct on epifluorescence images.

#### Cell cycle quantification

Cell cycle duration was quantified from septation to septation event. On transmitted light movies containing the entire transition from yeast to hypha we started our quantification by recording at what time the hypha septated at the end of the movie and then “went back in time” to the beginning of the movie following the lineage of the selected hypha and recording the time of each septation events in the lineage from final hypha to initial yeast. Results were averaged by cell type and over three experiments. Box plots were generated with http://shiny.chemgrid.org/boxplotr/

#### Nuclear and septum positioning

To assess nuclear positioning in the cells we measured the lengths from each cell tip to the middle of the nucleus on cells expressing NLS-GFP and plotted both lengths as a ratio. For the induced forms we plotted growing end length over non-growing end length. Cells were averaged by cell type and over three experiments. To assess nuclear positioning in the cytoplasm in yeast we used the same data as for nuclear positioning in the cell as the entire cell length is filled with cytoplasm. To assess nuclear positioning in the cytoplasm in hyphae we calculated the lengths from the growing cell tip to the middle of the nucleus and from the middle of the vacuolization zone to the middle of the nuclei. The vacuolization zone is the region in front of the large vacuole where small vacuoles are continuously delivered to the large one. We plotted the growing end length over non-growing end length. Results were averaged over three experiments. To assess septum positioning we plotted a ratio of the length from the growing cell tips to the septum over the length of the cytoplasm from transmitted light images. We averaged by cell type and over two experiments.

#### Quantification of microtubules plus-end localization

To measure microtubule plus end positions in hyphae, we used the center of the nucleus as reference point and measured the distance to each microtubule plus end. 10 hyphae, with a total of 50 microtubules were quantified. Microtubules pointing towards the growing tip have a positive distance value; those growing into the vacuolar compartment have a negative value. These distances were plotted on a graph, shown in Figure 7E. The accompanying schematic drawing indicates interval distances for the position of the vacuole and the two cell ends.

#### Microtubule depolymerizing drug washing

In microfluidic chambers we flowed cells with EMM-ALU supplemented with MBC (25µg/ml final concentration) for three hours before washing with EMM-ALU only. We assessed nuclear positioning in a strain expressing NLS-GFP before and after the wash. Even though microtubule re-polymerization occurred in the first 10 minutes (data not shown), we quantified the nuclear centering 50 minutes after wash because some cells were slower to reposition their nuclei than others.

**Table S1: Strains used in this study**

**Movie S1. Yeast-to-hypha transition of *S***. ***japonicus***. DIC movie of a growing *S. japonicus* mini colony in a microfluidic chamber in presence of the inducer RGE. Time is in h:min. Scale bar: 5µm.

**Movie S2. Tea1 deposition at hyphal tips.** Spinning disk movie of the hyphal form tagged with Tea1-GFP. Images are maximum intensity projections of 11 z-stacks (0.5µm step size). Time is in min:sec. Scale bar: 5μm.

**Movie S3. Vesicle trafficking in the yeast form.** Spinning disk movie showing middle plane section of the yeast form tagged with GFP-Ypt3. Time is in min:sec. Scale bar: 5µm.

**Movie S4. Vesicle trafficking in the hyphal form.** Spinning disk movie showing middle plane section of the hyphal form tagged with GFP-Ypt3. Time is in min:sec. Scale bar: 5µm.

**Movie S5. Microtubule organization in the yeast form.** Spinning disk movie of the yeast form tagged with GFP-Atb2. Images are maximum intensity projections of 10 z-stacks (0.5µm step size). Time is in min:sec. Scale bar: 5µm.

**Movie S6. Microtubule organization in the hyphal form.** Spinning disk movie of the hyphal form tagged with mCherry-Atb2. Images are maximum intensity projections of 9 z-stacks (0.5µm step size). Time is in min:sec. Scale bar: 5µm.

**Movie S7. Asymmetrical division in the hyphal form.** DIC movie showing hyphal growth and division. The front daughter cell rebuilds a vacuole after division. Time is in h:min. Scale bar: 5µm.

**Movie S8. Vacuole fusion.** DIC movie showing fusion of smaller vacuoles into an increasingly larger one. Time is in h:min. Scale bar: 5µm.

## Supporting information

## Acknowledgements

We thank Snezhana Oliferenka (Crick Institute, London) for critical technical help at the start of the project. We thank her and Hironori Niko (National Institute of Genetics, Japan) for strains and reagents. This work was supported by an ITN funding (FungiBrain) and SNF grant (310030B_176396) to SGM. We are thankful to Serge Pelet and his group, as well as the Martin group for critical reading of the manuscript.

## Author contributions

Conceptualization: CK and SGM. Investigation: OD discovered the inducing action of fruit extracts and constructed the Pom1-GFP strain; CK performed all experiments. Writing: CK wrote the original draft; SGM and CK edited it. Funding acquisition: SGM.

---

**Figure S1.**
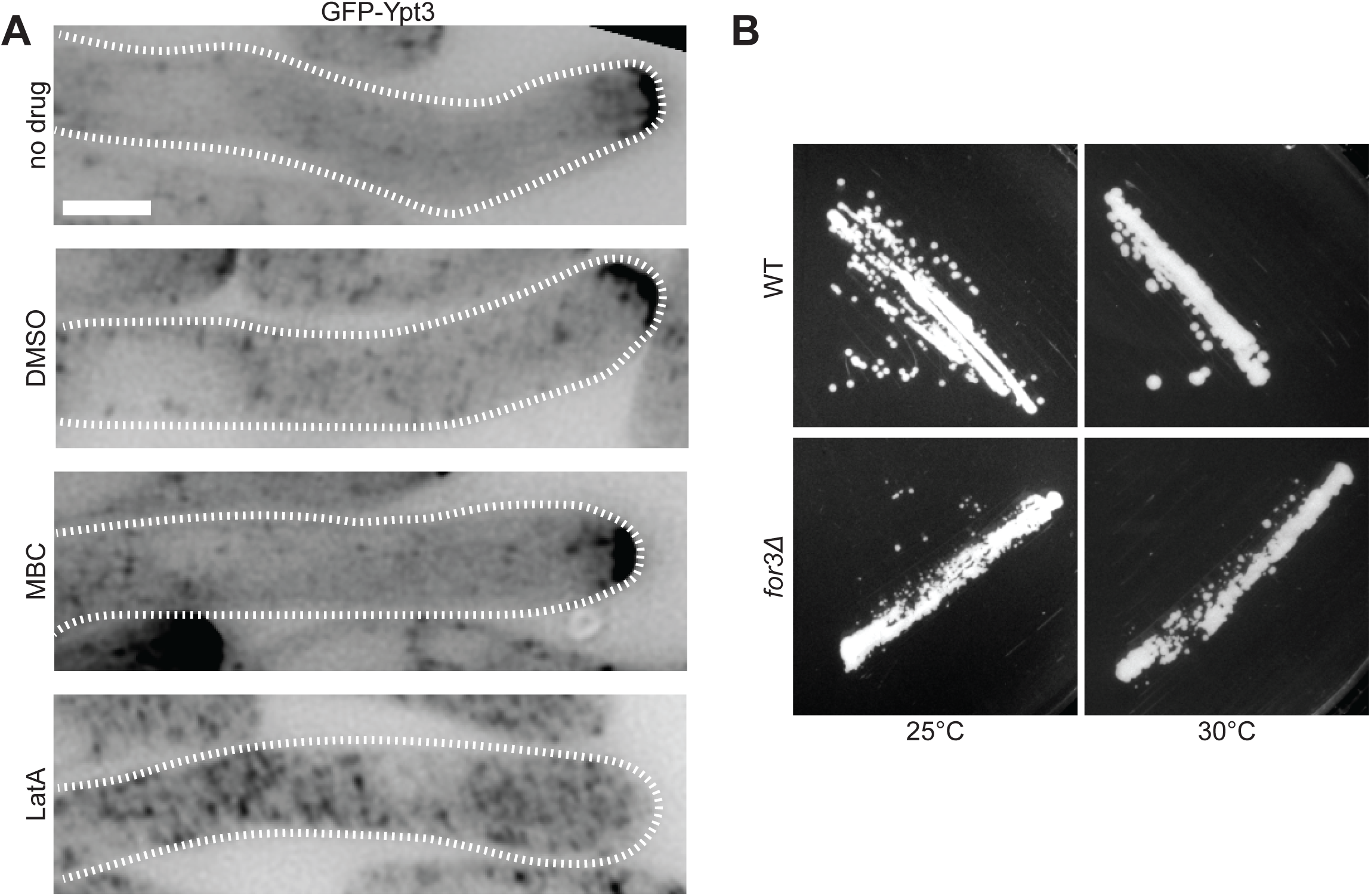
Importance of formin For3 and F-actin in polarized growth. **A**. Middle plane images of GFP-Ypt3 in hyphae grown in a microfluidics chamber in presence or not of DMSO, MBC or LatA. **B.** *S. japonicus* WT and *for3Δ* strains growing on solid rich media for three days at 25°C or 30°C. Scale bar: 5µm.

**Figure S2.**
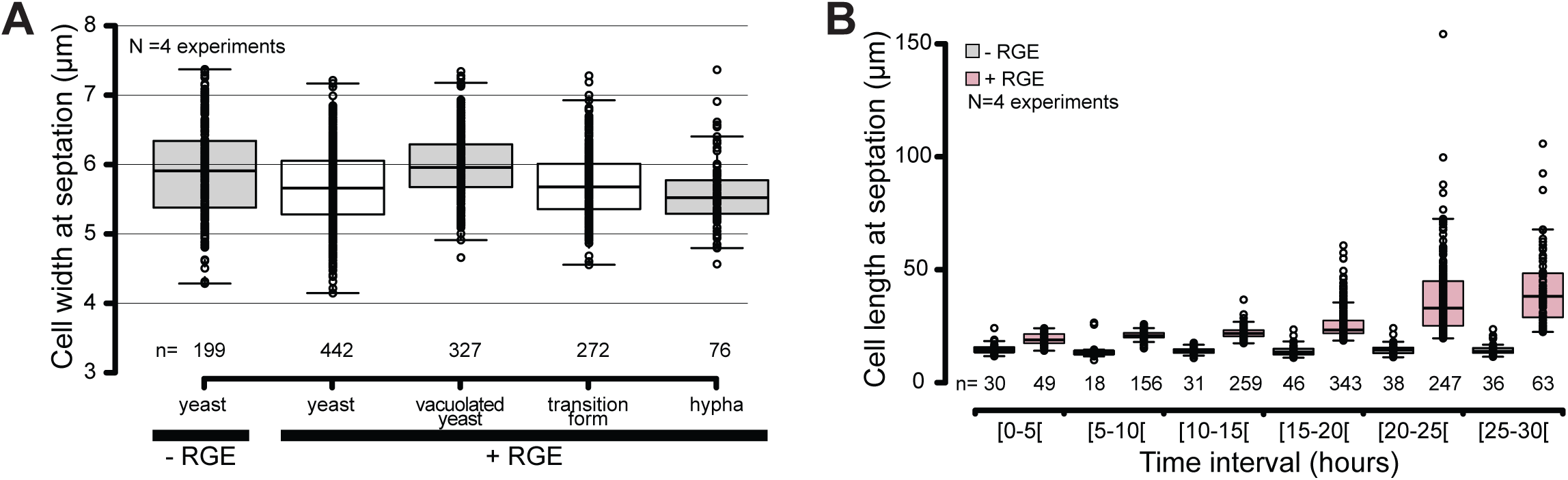
Measurement of cell width and length at septation. **A**. Box plot of cell width at septation in the different morphological forms of *S. japonicus*. **B**. Box plot of septation length of cell populations over 30 hours in microfluidic chambers. Box plots show first and third quartile and median, whiskers extend times the interquartile range from the first and third quartile.

**Figure S3.**
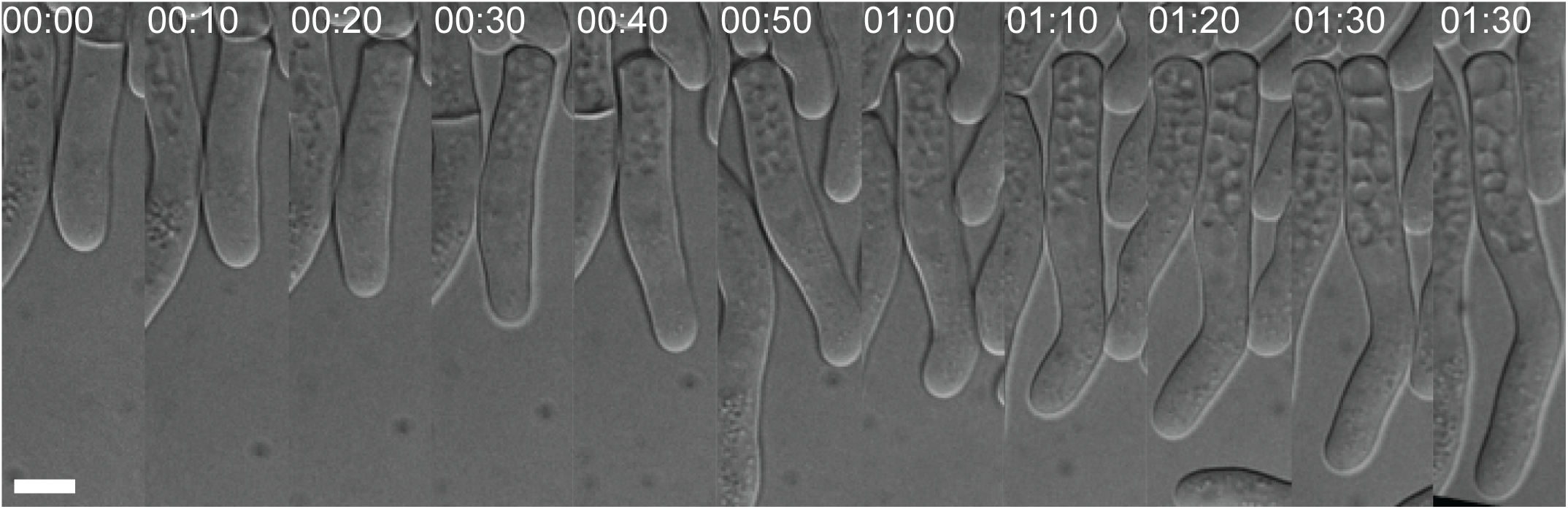
Asymmetric formation and partitioning of vacuoles in the transition form. DIC microscopy image showing division of a cell growing in RGE highlighting vacuole formation at the back of the front daughter cell. Time in h:min. Scale bar: 5µm.

**Figure S4.**
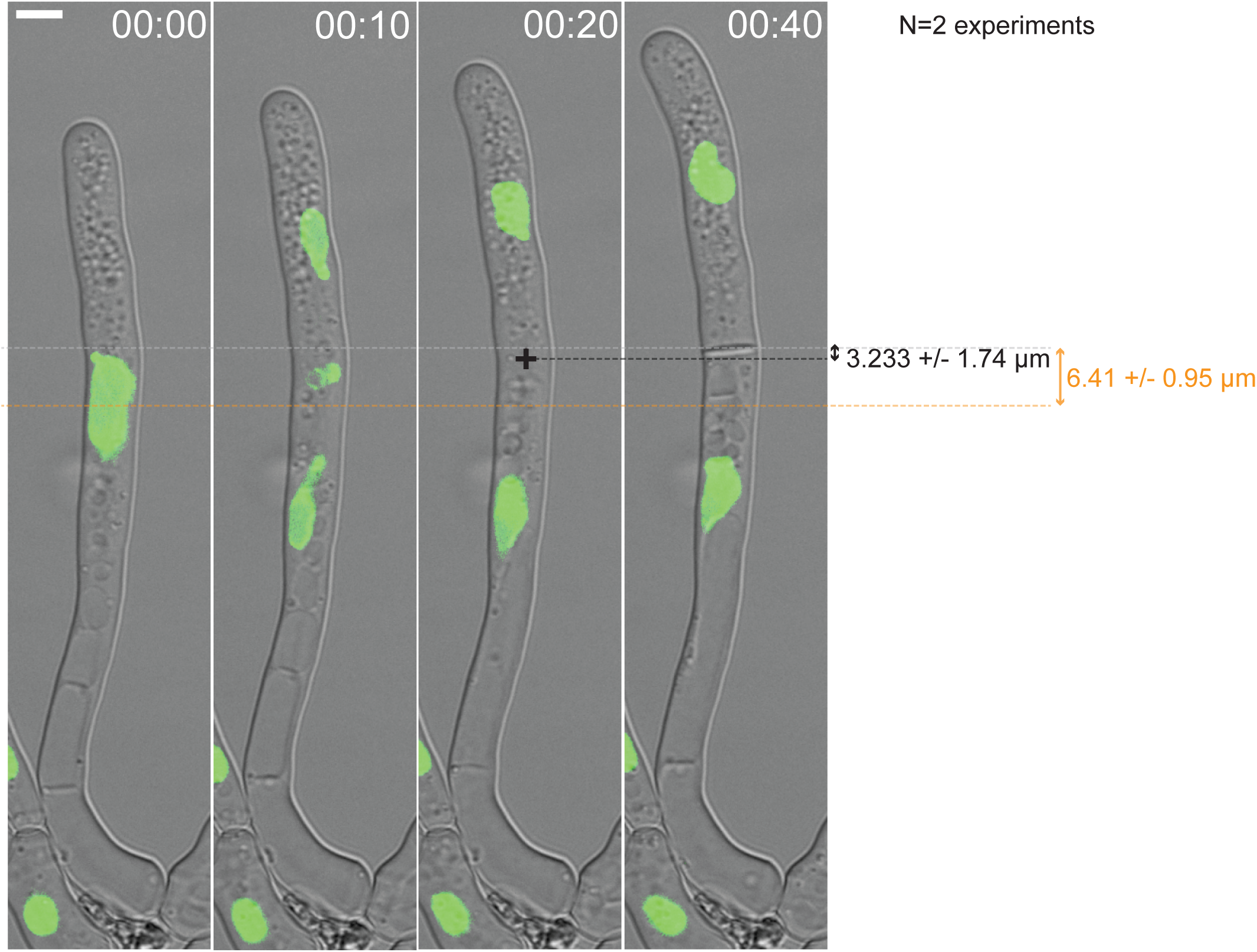
Septum position is better predicted by the middle of the anaphase spindle than the position of the pre-divisional nucleus. Timelapse imaging of a hypha where the positions of the pre-divisional nucleus (orange dotted line), the inferred middle of the anaphase spindle (black cross) and the septum (grey dotted line) are marked. The arrows show the distance between the position of the septum and that of either the pre-divisional nucleus or the middle of the anaphase spindle with their respective average and standard deviations over 20 cells. Time in h:min. Scale bar: 5µm.

